# Deconstructing multivariate decoding for the study of brain function

**DOI:** 10.1101/158493

**Authors:** Martin N. Hebart, Chris I. Baker

## Abstract

Multivariate decoding methods were developed originally as tools to enable accurate predictions in real-world applications. The realization that these methods can also be employed to study brain function has led to their widespread adoption in the neurosciences. However, prior to the rise of multivariate decoding, the study of brain function was firmly embedded in a statistical philosophy grounded on univariate methods of data analysis. In this way, multivariate decoding for brain interpretation grew out of two established frameworks: multivariate decoding for predictions in real-world applications, and classical univariate analysis based on the study and interpretation of brain activation. We argue that this led to two confusions, one reflecting a mixture of multivariate decoding for prediction or interpretation, and the other a mixture of the conceptual and statistical philosophies underlying multivariate decoding and classical univariate analysis. Here we attempt to systematically disambiguate multivariate decoding for the study of brain function from the frameworks it grew out of. After elaborating these confusions and their consequences, we describe six, often unappreciated, differences between classical univariate analysis and multivariate decoding. We then focus on how the common interpretation of what is signal and noise changes in multivariate decoding. Finally, we use four examples to illustrate where these confusions may impact the interpretation of neuroimaging data. We conclude with a discussion of potential strategies to help resolve these confusions in interpreting multivariate decoding results, including the potential departure from multivariate decoding methods for the study of brain function.

**Highlights:**

- We highlight two sources of confusion that affect the interpretation of multivariate decoding results
- One confusion arises from the dual use of multivariate decoding for predictions in real-world applications and for interpretation in terms of brain function
- The other confusion arises from the different statistical and conceptual frameworks underlying classical univariate analysis to multivariate decoding
- We highlight six differences between classical univariate analysis and multivariate decoding and differences in the interpretation of signal and noise
- These confusions are illustrated in four examples revealing assumptions and limitations of multivariate decoding for interpretation

## 1. Introduction

Multivariate decoding^1^ has become a central method for the analysis of neuroscientific data. It is being employed commonly in fMRI (Haynes, 2015; Haynes and Rees, 2006; Norman et al., 2006; Tong and Pratte, 2012), but also neurophysiology in non-human primates (Quian Quiroga and Panzeri, 2009) and humans (Contini et al., 2017). The approach grew rapidly in popularity in the neuroimaging community when it became clear that it was not only useful for classification related to real-world applications such as brain-computer interfaces, but also for studying brain function. Now, in many domains classical univariate methods have been replaced by multivariate decoding, in part owing to the higher sensitivity afforded by these techniques (Haynes and Rees, 2006; Norman et al., 2006). In this way, multivariate decoding for brain interpretation grew out two established approaches: multivariate decoding for predictions in real-world applications, and classical univariate analysis for the study of brain function.

In this article, we argue that rather than being part of a consistent and independent statistical framework, multivariate decoding for brain interpretation often reflects a mixture of the philosophies it originated from (Figure 1A), one activation-based and the other information-based. As a consequence, this mixture of philosophies creates a lot of potential for confusion in the interpretation of results derived from multivariate decoding methods. The aim of this article is to provide a systematic understanding of multivariate decoding for the study of brain function and the assumptions and limitations of this approach in the interpretation of multivariate decoding results.

**Figure 1.**
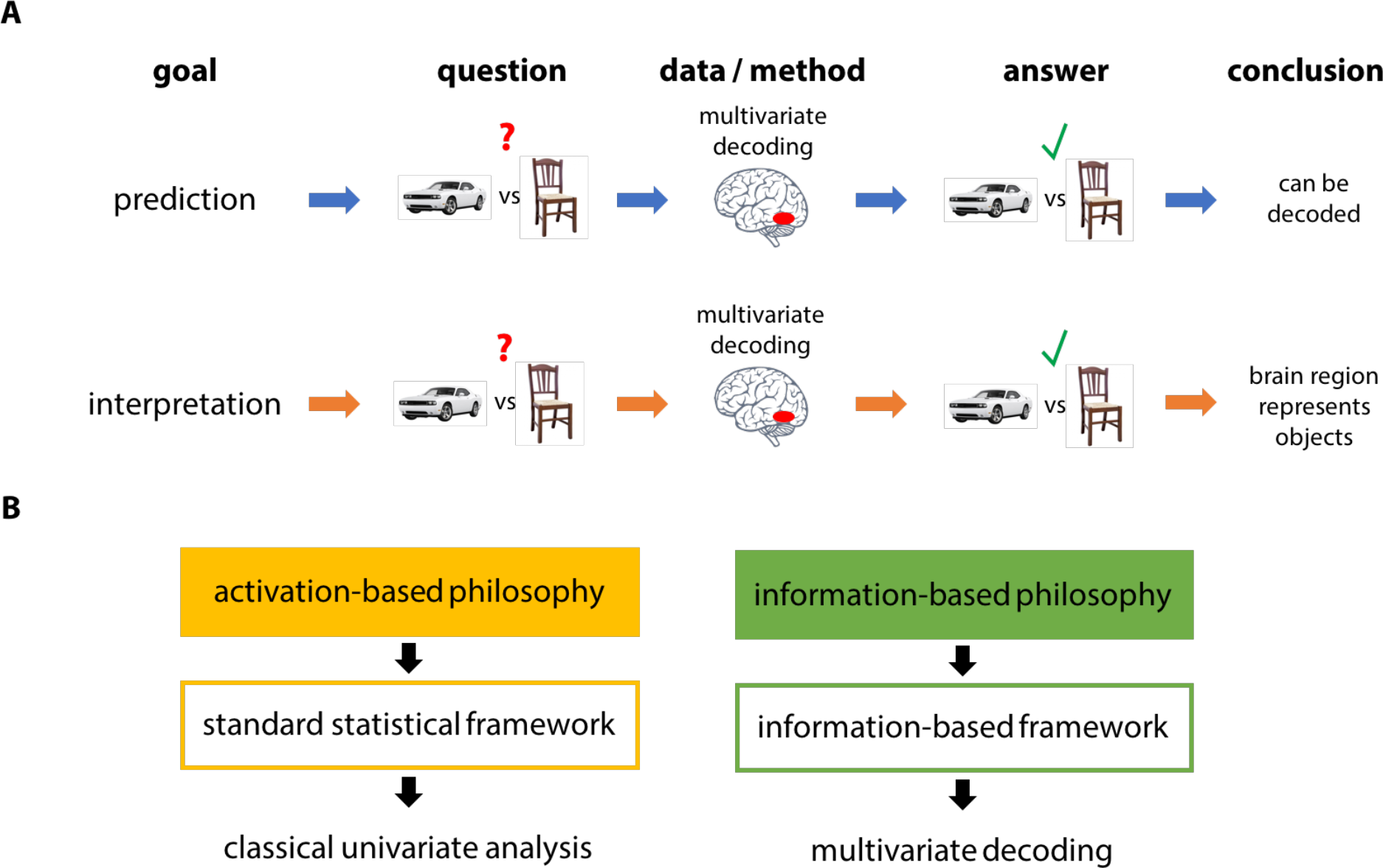
The two sources of confusion in multivariate decoding. A. Multivariate decoding was developed for predictions in real-world applications, but is widely used for interpretations about brain function. Since both approaches are often treated as a unitary method despite making different assumptions, this provides a source for confusion. B. The choice between classical univariate analysis is not only a choice of method but a choice of underlying philosophy, activation-based or information-based. Confusion can arise when the conceptual and statistical framework underlying classical univariate analysis is applied to multivariate decoding.

First, we describe the two sources of confusion: i) the mixture of multivariate decoding for prediction and multivariate decoding for interpretation, and ii) the mixture of the statistical and conceptual philosophies underlying classical univariate analysis and multivariate decoding. Next, we illustrate six methodological and interpretational changes that – explicitly or implicitly – are adopted when shifting from classical univariate methods to multivariate decoding. This discussion is important, because it shows how multifaceted the differences between these approaches are and why they have been so difficult to characterize. Moving to a purely multivariate description of data, we then describe how the meaning of signal and noise is different in the statistical frameworks underlying classical univariate analysis and multivariate decoding. Finally, using four illustrative examples we demonstrate how the sources of confusion can affect the interpretation of multivariate decoding results.

Throughout the article, we use functional MRI as an example, where multivariate data are multiple voxels measured at different time points, and where predicted variables are experimental conditions^2^. However, this discussion applies equally to other modalities (e.g. structural MRI, MEG/EEG, connectivity measures) whenever multivariate decoding is used as a method of data analysis. In addition, we focus our discussion of multivariate decoding on multivariate classification, although our arguments may apply equally to multivariate regression in a decoding setting.

## 2. Two sources of confusion

### Multivariate decoding for prediction vs. interpretation

The first major source of confusion stems from the distinction between multivariate decoding for prediction and multivariate decoding for interpreting brain function (Figure 1A), which can be illustrated by the results of the 2006 Pittsburgh Brain Activity Interpretation Competition. The purpose of the competition was to use brain activity data measured with fMRI to predict the subjective perception of movie segments according to several criteria including the objects, spatial locations, sounds, and emotions associated with these segments. The winner was determined by who best predicted ratings based on independent fMRI data. According to the competition website and call for submissions, the goals of the competition were “to advance the methodology and assess the state of the science”, and “to advance the understanding of how the brain encodes, represents, and operates on dynamic experience”^3^. The competition received a lot of interest in the community, with multiple participants using multivariate decoding methods including sophisticated machine learning algorithms to carry out predictions (Nature Neuroscience Editorial, 2006). Surprisingly, the winners of the contest were a team of data scientists who acknowledged they did not know much about the brain prior to the competition (Sona et al., 2007). When visualizing the voxels their classifier used for predictions, many of them were contained within the ventricles and other regions typically related to physiological noise. Potentially, the most predictive voxels did not reflect brain activity in response to the ratings, but rather head motion and changes in physiological noise. Thus, one important lesson learned through the competition in 2006 is that the use of multivariate decoding can lead to excellent predictions, but sometimes to not very useful interpretations in terms of brain function. Perhaps for this reason, in 2007 the competition included a separate neuroscience prize for making substantial contributions to the understanding of brain function.

Today, the dichotomy of maximal prediction on the one hand and interpretation of brain function on the other continues to be of importance^4^. *Multivariate decoding for prediction* aims at identifying biomarkers that can be used to carry out predictions about underlying states of the brain. Here, maximal decoding performance is the goal, and success is determined by a model that can decode mental or physiological states from previously unseen data with high accuracy. The most frequently used tools in multivariate decoding are machine learning classifiers or variants thereof, which are often treated as a black box approach to assign labels to available data. Among others, studies employing multivariate decoding for prediction have investigated the prediction of disease status and progression (Ewers et al., 2011; Orrù et al., 2012), the usefulness of neuroimaging for brain computer interfaces in quadriplegic patients (Blankertz et al., 2007), and the feasibility of neuroimaging-based lie detection (Davatzikos et al., 2005; Farah et al., 2014; Peth et al., 2015). In addition, multivariate decoding for prediction has been used for read-out of information from visual cortex during perception (Kay et al., 2008; Miyawaki et al., 2008; Naselaris et al., 2009; Nishimoto et al., 2011; Thirion et al., 2006) and during sleep (Horikawa et al., 2013), and from auditory cortex during speech (Formisano et al., 2008). The source of the information is not necessarily of interest to these approaches, as long as the prediction is successful and can generalize to other relevant datasets^5^.

In contrast, *multivariate decoding for interpretation* aims at a better understanding of the human brain and does not require high predictive accuracy. The reasoning behind this approach is that as soon as a decoding model performs reliably better than chance, this demonstrates that there is structure in the data with respect to the conditions of interest, for example whether the participant was presented with a picture of a car or a chair. From this the researcher typically concludes that a given brain region carries discriminative information^6^ about these categories, which may enlighten us about the neural computations carried out in this brain region. Among others, multivariate decoding for interpretation revealed the existence of subcortical effects of binocular rivalry (Haynes et al., 2005), feature binding in primary visual cortex (Seymour et al., 2009), working memory representations in primary visual cortex (Harrison and Tong, 2009), unconscious intentions in frontopolar cortex (Soon et al., 2008), visual search templates in object-selective cortex (Peelen et al., 2009), and reward value representations in parietal cortex (Kahnt et al., 2014). For this approach, variables such as head motion would act as confounds even when they consistently co-occur with the experimental variables.

While this distinction between prediction and interpretation was made explicit early on (Norman et al., 2006), multivariate decoding is commonly being treated as one methodological entity that can be applied equally for both approaches (for review, see Tong and Pratte, 2012). What has often been overlooked, however, is that the tools of multivariate decoding – machine learning algorithms – were not developed for the interpretation of brain function, but simply for making predictions about variables based on available data. In the context of the interpretation of brain imaging results this has two consequences: i) any interpretation that goes beyond the existence of a statistical dependence, i.e. beyond the presence of information about experimental variables in brain imaging data, may come with additional assumptions that might be violated and may invalidate this interpretation; ii) the limitations imposed by multivariate decoding for prediction may unnecessarily constrain the use of multivariate decoding methods in the context of interpretation^7^. While both consequences deserve study, most of this article will focus on the first of these two: the interpretation of brain imaging data that goes beyond the presence of information.

### The statistical frameworks underlying classical univariate analysis and multivariate decoding

The second major source of confusion concerns differences in the conceptual and statistical philosophies underlying classical univariate analysis and multivariate decoding (Figure 1B). Classical univariate analysis and multivariate decoding are much more than just methods of data analysis. They are embedded in separate philosophies about the nature of neuronal representations, one activation-based, and the other information-based. These philosophies are manifested in different statistical frameworks. In this sense, classical univariate analysis is an approach to study brain activation *within* a standard statistical framework, while multivariate decoding is an approach to study information-content *within* an information-based framework. The exact implementation of each approach, for example the use of a general linear model (GLM) in univariate analysis or a linear classifier in multivariate decoding, carries assumptions specific to these frameworks.

The *activation-based philosophy* has been the dominant thinking in the interpretation of neuroscientific results. It is based largely on the analysis of different levels of brain activity. In this view, a higher firing rate of a neuron is interpreted as a stronger engagement of that neuron in the process of study^8^. The same reasoning is applied in other domains, such as a larger BOLD response in an MRI voxel, increased voltage deflections in an EEG channel, or power increases in frequency bands of MEG. Analysis of brain structure or connectivity follows a similar scheme, where their relevance to the process of study is determined by changes in relation to an experimental variable. Importantly, this activation-based philosophy is not limited to univariate analysis, but can be extended to multivariate analysis, when a pattern of conjoint activation is the focus of study. This philosophy, however, does not underlie the statistical framework of multivariate decoding. Instead, multivariate decoding is embedded in an *information-based philosophy*, which focuses on the information contained in a brain region and how this information may be communicated to other parts of the brain. Here, *any* measurable difference between the conditions of interest, or more precisely mutual information between experimental variables and brain data, can be interpreted as reflecting the process of study (Kriegeskorte and Bandettini, 2007). How these differences in philosophy affect our interpretation of brain responses, however, has been largely ignored^9^.

Importantly, each of these philosophies has been associated with a statistical framework that formalizes the assumptions of the philosophy, allowing estimation of the relevant quantities (activation vs. information), and providing statistical tests to evaluate the generalizability of these estimates. The activation-based philosophy commonly uses a *standard statistical framework*, which reflects both the statistical model underlying most activation-based analyses and the chosen paradigm for statistical inference. The dominant statistical paradigm in the standard statistical framework is classical frequentist statistics, although Bayesian statistics can also be used for statistical inference. A very common feature in the standard statistical framework is the use of a linear model that tests for a linear relationship between model variables and measured data, and statistical inferences are typically carried out on the estimates derived from this model (e.g. a t-test on an estimate of the mean).

In contrast, the information-based philosophy relies on an *information-based framework* derived from information theory, in which statistical estimation is carried out using mutual information or related measures. While the standard statistical framework is typically limited to testing a specific – mostly linear or monotonic – relationship between data and experimental variables, the information-based framework relies on *any* differences in data distributions between pairs of variables, including nonlinear as well as non-monotonic effects. In that sense, the information-based framework is more general than the standard statistical framework^10^. Instead of directly estimating mutual information, which has been very difficult with limited data (but see Ince et al., 2017), other statistical analyses that derive information estimates can be used. From a statistical point of view, multivariate decoding is one such analysis, and classification accuracy is one form of information estimate. Importantly, since multivariate decoding does not provide a framework for inferential statistics, the statistical analysis of decoding results usually borrows methods from other statistical inference paradigms.

Here we argue that the current thinking in multivariate decoding in the interpretation framework is not information-based, but still largely embedded in i) an activation-based philosophy that was adopted from classical univariate analysis and ii) the standard statistical framework including the statistical model underlying most univariate analysis. As will become clear, this mixture can lead to non-intuitive interpretations of what is considered signal and noise in a multivariate pattern. In addition, it leaves us with a mixture of the analysis repertoire from activation-based analysis and multivariate decoding, and provides the potential for confusion.

## 3. Differences between classical univariate analysis and multivariate decoding

Commonly, the use of multivariate decoding over univariate analysis is justified by two factors: i) the increased sensitivity in detecting meaningful differences in the brain by combining information across multiple voxels (Haynes and Rees, 2006; Norman et al., 2006, but see Allefeld et al., 2016) and ii) the increased specificity in being able to access widely distributed population codes by the joint analysis of multiple voxels that would not be available by assessing each voxel separately (Haynes, 2015; Kriegeskorte, 2011)^11^. While both factors describe the motivation for using multivariate analysis, it is important to realize that there are multiple changes that are a consequence of this departure from classical univariate analysis. In the following, we highlight six specific changes and illustrate the reasons for these changes (Figure 2). While there is some overlap between these changes and while some of the changes are prerequisites of others, none of them necessarily co-occur, i.e. they can be treated as largely independent. Consequently, this allows us to pinpoint the changes that are truly necessary for the increase in sensitivity and specificity, and those that are a mere reflection of the specific method of choice.

**Figure 2.**
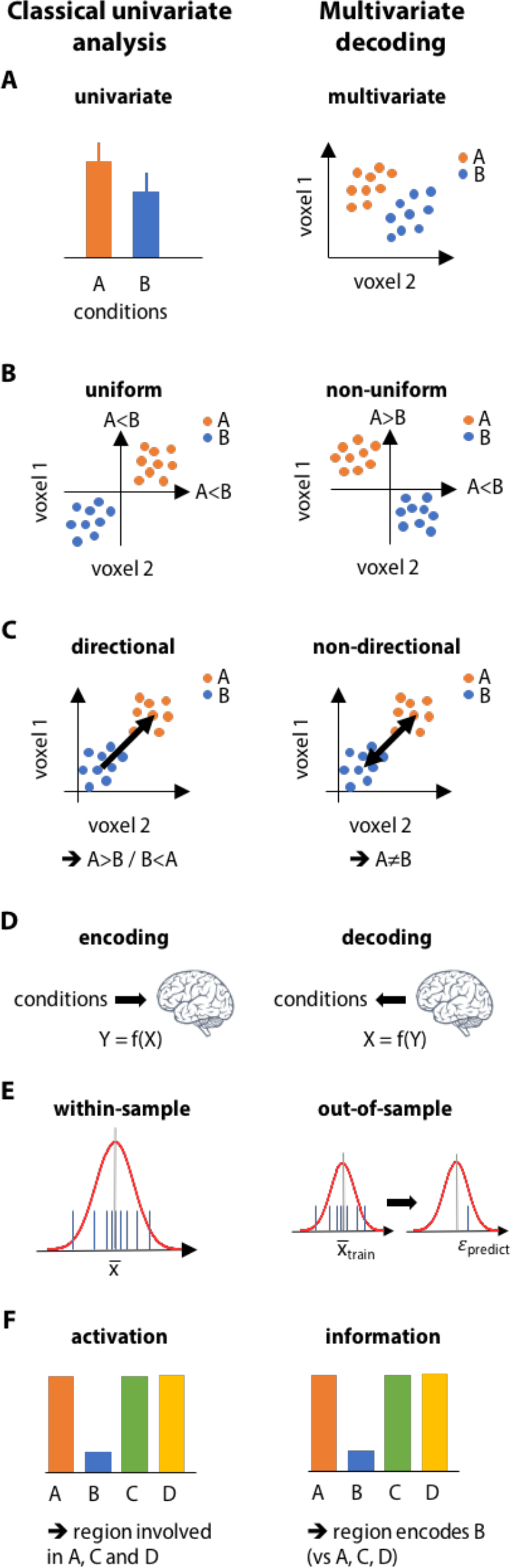
*Six differences between classical univariate analysis and multivariate decoding.*

### 1. Univariate vs. multivariate

The most obvious difference between the two approaches is already part of their respective names and denotes the difference between univariate and multivariate analysis (Figure 2A). While univariate analysis refers to a separate analysis of each individual voxel, multivariate analysis refers to the joint analysis of multiple voxels^12^. In classical univariate analysis, voxels are typically only combined by pooling measurements within predefined regions of interest or by applying spatial smoothing. However, this approach largely ignores the *relevance* of each voxel in distinguishing between experimental conditions and does not utilize the covariance between voxels. In contrast to univariate analysis, multivariate analysis allows optimally combining voxels by taking into account each voxel’s contribution to discriminability. In addition, the covariance between voxels carries additional information that can be exploited in multivariate analysis.

### 2. Uniform vs. non-uniform response sign

In classical univariate analysis, regions-of-interest are typically described by a set of neighboring voxels that exhibit relatively uniform responses. The voxels may fluctuate in the response level, but are assumed to be of the same sign, and within regions these differences are typically not interpreted. For example, while it is known that different voxels in the fusiform face area (FFA) respond to faces to different degrees, it is nevertheless assumed that FFA has a uniform, positive response sign to faces.

In multivariate decoding, voxels in a region can show non-uniform response signs: Both activation and deactivation in neighboring voxels is interpreted as being informative about the variable of interest, and both signs contribute to the overall estimate of information content (Figure 2C, right). In other words, in multivariate decoding it is not important that all voxels of a brain region show responses of the same sign; positive and negative responses are equally meaningful. To clarify, by non-uniform we are not referring just to any variations in responses between neighboring voxels, which would be a property of what we described as “multivariate” above; rather, we specifically refer to the fact that one voxel can show a positive response while the neighboring voxel can show a negative response. Indeed, it is possible to restrict a multivariate analysis to uniform responses, although in many cases this requires the development of new methods of data analysis or an adaptation of existing methods (e.g. Hebart et al., 2014b).

### 3. Directional vs. non-directional analysis

In classical univariate analysis, a brain region is said to be engaged in a cognitive process when it responds more to the experimental condition than a control condition, or when it shows an overall positive or negative relationship with different levels of the experimental variable. The same contrast is calculated for each voxel individually, and overall it is determined whether a brain region is activated or deactivated (Figure 2C, left). Estimates of activation or deactivation can then be taken from the subject to the group level, and additional statistical analysis can be used to infer whether the population exhibits activation or deactivation in that brain region. This describes a *directional* analysis, because the sign of the difference is taken to be important (more activated or more deactivated than control). While non-directional analyses (e.g. F-tests) are possible in classical univariate analysis, they are much less common and are usually not employed to draw inferences at the subject level.

In multivariate decoding, an analysis is almost always carried out in a *non-directional* manner. This is not surprising, because in a multivariate space direction does not have much of a meaning. For example, one voxel may be more activated in one condition than another, while another voxel may be less activated. This makes it impossible to describe a response direction as overall positive or overall negative and thus makes it hard to assign meaning to this “mixture in directions”. For most analyses, the direction does not matter anyway, because the focus lies on the discriminability between patterns of activity and not the difference between individual voxels^13^.

It is, however, possible to carry out a directional analysis in multivariate decoding, and there are at least two cases where directional analysis may make sense in the context of multivariate analysis. First, when there are uniform response differences as described above, a directional multivariate analysis describes a direction in voxel space that is related to the general activation or deactivation of a region. This multivariate analysis would be more sensitive than a classical univariate analysis, because it would allow optimally combining voxels across the region. Second, even for non-uniform response differences if the assumption is that the difference in response patterns between conditions is reproducible across subjects, then the direction indeed matters and is required to draw inferences at the population level about “representative” response differences. Indeed, it has been suggested that those differences can be analyzed at the group level in a directional manner (Gilron et al., 2017). In contrast, if the focus lies merely on the discriminability of patterns, then a non-directional analysis is ideal. To sum up, both directional and non-directional analyses can be meaningful in multivariate decoding, and non-directional analysis is not a necessary aspect of multivariate decoding.

### 4. Encoding vs. decoding

Encoding describes the prediction of data (dependent variables) from experimental conditions (independent variables), whereas decoding describes the prediction of experimental conditions from data (Figure 2B). For example, a GLM in a classical univariate analysis is an encoding model, because it provides a (high-level) description of how a process of study is encoded in a brain response^14^. It has been argued repeatedly that encoding and decoding are complementary when the goal is to quantify a statistical dependence between dependent and independent variables (Friston et al., 2008; Kriegeskorte, 2011; Naselaris et al., 2011). Decoding is commonly used in multivariate data analysis not because it offers a computational benefit over encoding, but because of its apparent simplicity, appeal, and novelty. Decoding analyses are relatively easy to carry out, for example with out-of-the-box classification algorithms (e.g. as implemented in LIBSVM, Chang and Lin, 2011), or by using the popular correlation-based classifier that requires only the computation of a small number of correlations across voxels (Haxby et al., 2001). Part of the appeal of decoding came from the idea that decoding may have access to fine-scale information beyond the resolution of fMRI (Kamitani and Tong, 2005, but see Freeman et al., 2011; Op de Beeck, 2010) and the possibility to describe these methods as tools for “mind-reading” (Haynes and Rees, 2006; Norman et al., 2006). In addition, some treat an activity pattern as an explicit representation of the variable of interest, and thus linear decoding may be used to describe what information about this represented variable can be “read out” by other parts of the brain (Diedrichsen and Kriegeskorte, 2017; Kriegeskorte, 2011). However, decoding also has downsides. In contrast to encoding, it does not allow a complete functional description of brain regions (Naselaris et al., 2011). In addition, with decoding it is not possible to calculate “noise ceilings” to determine whether limitations in characterizing a statistical dependence are related to the model or the data quality (Naselaris et al., 2011).

It is worth noting that multivariate *encoding* approaches with similar potential to multivariate *decoding* have been suggested previously, such as MANCOVA (Friston et al., 1995), canonical correlation analysis (Friman et al., 2001) or partial least squares (McIntosh and Lobaugh, 2004). However, they have not received as much attention as multivariate decoding or have been used to address different questions. There are multiple reasons for this discrepancy, including interpretational complexity, problems arising from fitting a model with more parameters than measurements (“curse of dimensionality”), the inability to generate unbiased estimates that could easily be translated from the subject level to the group level (Allefeld and Haynes, 2014; Walther et al., 2016), or distributional assumptions (Kriegeskorte, 2011; Kriegeskorte and Diedrichsen, 2016). In contrast, multivariate decoding promises a gain in sensitivity while avoiding these particular issues.

### 5. Within-sample statistical estimation vs. out-of-sample prediction

Classical univariate analysis relies on the use of *within-sample statistical estimation* (Figure 2E, left). In this approach, all available data are first used to attain statistical estimates of how the experimental variables map to the data (e.g. beta weights in a GLM estimated on fMRI data). Then, those “activation estimates” are subjected to statistical tests (e.g. *t*-tests) to determine whether the results would generalize to the population. In multivariate decoding, the goal is not to attain activation estimates, but estimates of the information about experimental variables contained in the data. An estimate of information content can be quantified as the predictive value of a model using *out-of-sample prediction* (Figure 2E, right). In out-of-sample prediction, a researcher first estimates a model on a subset of the available data and then uses this model to predict the experimental variable associated with the left-out data^15^. In multivariate decoding, this prediction is typically quantified in terms of classification accuracy, correlation coefficient, or explained variance. When this process of model estimation and out-of-sample prediction is carried out iteratively on different subsets of the data, this approach is described as cross-validation. Importantly, out-of-sample prediction still requires a statistical test to determine whether a given estimate of information content (e.g. classification accuracy) is reliable, even when the prediction is very good (Combrisson and Jerbi, 2015; Isaksson et al., 2008). Statistical testing procedures on cross-validated information estimates require additional scrutiny (Görgen et al., this issue; Jamalabadi et al., 2016; Noirhomme et al., 2014; Schreiber and Krekelberg, 2013). Thus, the crucial difference lies not in the statistical procedure (e.g. Bzdok and Yeo, 2017), but in the approach for achieving (unbiased) estimates of the variables of interest, for example activation means in classical univariate analysis or classification accuracies as estimates of information content in multivariate decoding. In that respect, the term “out-of-sample estimation” may in some cases be more telling than “out-of-sample prediction”.

Out-of-sample prediction is the typical approach in multivariate decoding, because in most cases multivariate models have many more degrees of freedom than univariate models and can much more easily overfit to the idiosyncrasies of the data, leaving us with biased estimates of information content (Bzdok, 2016). Multivariate methods such as MANOVA or pattern component modeling (Diedrichsen et al., 2011; this issue), which do not require out-of-sample prediction, can reduce this complexity with additional assumptions about the distribution of the data. However, while growing in popularity, such methods are not commonly employed. As discussed above, there may be doubt that the assumptions of those multivariate tests hold for fMRI data in practice, while out-of-sample prediction does not require those assumptions (Kriegeskorte, 2015). However, as an alternative to more traditional statistical tests, procedures such as permutation tests can be used to carry out within-sample estimation even for multivariate decoding, without requiring cross-validation (Kriegeskorte et al., 2006).

### 6. Activation vs. information

As pointed out above, classical univariate analysis and multivariate decoding are embedded in activation-based and information-based philosophies, respectively (Figure 2D; Kriegeskorte and Bandettini, 2007). Take an imaginary region that responds to faces and not to objects. According to the activation-based view, this region would be described as face-selective. However, now assume the region additionally responds to gratings, scrambled objects, and even when nothing is presented. In other words, the region is always active and only becomes silent when an object is shown. While according to the activation-based view it would represent anything but objects, in the information-based view this region is maximally informative about the presence of objects (Figure 2D). This is because the inactivity and activity in both cases carry information about the presence or absence of an object (Panzeri et al., 2015). This example naturally extends to the multivariate analysis of voxels: A pattern of activity can represent many more different states than each voxel individually. The idea of a widely-distributed population code has motivated the study of multivariate patterns in terms of information content (Cox and Savoy, 2003; Haxby et al., 2001; Kay et al., 2008; Naselaris et al., 2009). Further, additional information may come from studying not only the mean response pattern, but also the variability (Averbeck et al., 2006; Panzeri et al., 2015). The information contained in the variability of response patterns will be discussed in more detail in the Section 3 (“What is signal and what is noise in multivariate decoding?”).

### What differences are necessary for increased sensitivity and specificity?

The fact there are at least six distinct differences between classical univariate analysis and multivariate decoding might explain why it has been so difficult to compare the two methodologies directly (Coutanche, 2013; Davis et al., 2014; Jimura and Poldrack, 2012; Smith et al., 2011). Returning to the original motivation that stimulated the shift towards multivariate decoding, it becomes clear that only two of these six differences are strictly necessary for a benefit over classical univariate analysis: increased sensitivity is achieved through the joint analysis of multiple voxels (*univariate* vs. *multivariate,* Figure 2A), and increased specificity through multivariate analysis in an information-based framework (*activation* vs. *information,* Figure 2D). The other four differences – uniform vs. non-uniform response signs, directional vs. non-directional analysis, encoding vs. decoding, and within sample estimation vs. out-of-sample prediction – are merely byproducts that may only be necessary for the specific methods that are commonly employed. For example, as mentioned earlier, multivariate analysis can be carried out separately for both uniform and non-uniform responses. Out-of-sample prediction on the other hand could – at least for some approaches – be replaced by appropriate permutation-based approaches^16^, which may even improve their sensitivity (Friston et al., 2008; Rosenblatt et al., 2016). But even within the two critical differences – multivariate analysis and the use of an information-based framework – it is worth discussing whether the focus should lie only on the *estimation* of response patterns and their distance and discriminability in a multivariate space, or whether *variability* of response patterns should also be treated as a meaningful source of information. This distinction will be covered in further detail in the following section.

## 4. What is signal and what is noise in multivariate decoding?

To appreciate how the differences between the activation-based and information-based philosophies described above affect our interpretation of brain signals, it is helpful to evaluate the differences in understanding of signal and noise in the standard statistical framework and the information-based framework, respectively.

### Signal and noise in the activation-based philosophy

In neuroscience, the measurement of a brain response is usually treated as a noisy observation of ground truth. Since we do not know what ground truth is, we can use a statistical model that allows us to formalize our assumptions about the brain response, in the hope this model provides a useful approximation of this ground truth. A popular choice for such a statistical model is a linear model that decomposes a measurement into different components. If weighted appropriately, those components would then provide a full description of the measured brain response. In classical parametric statistics, our goal is to estimate those weights or parameters based on our observations (e.g. beta weights in a GLM). This view reflects the activation-based philosophy, formalized through the standard statistical framework.

Figure 3A illustrates what is commonly perceived as signal and noise^17^, with the example of two experimental conditions depicted in orange and blue. Here, a signal reflects the *difference in the multivariate means* related to conditions of interest, represented as vectors in voxel space. Alternatively, the difference in multivariate means can be described as two multivariate patterns that are representative of those conditions of interest and that are different from each other. Noise is reflected in *error*, which describes the variability not accounted for by experimental conditions, and which can be either *condition-independent* or *condition-dependent* (Figure 3A, right). One noteworthy case of condition-dependent error are *confounds,* which are other variables that covary with the conditions of interest and which can influence their estimation. In a multivariate GLM, typical examples of an error component would be a condition-independent Gaussian with a given variance and covariance structure. Other, more complex generalized or hierarchical models could account for non-Gaussian error or condition-dependent error (e.g. heteroskedastic error).

**Figure 3.**
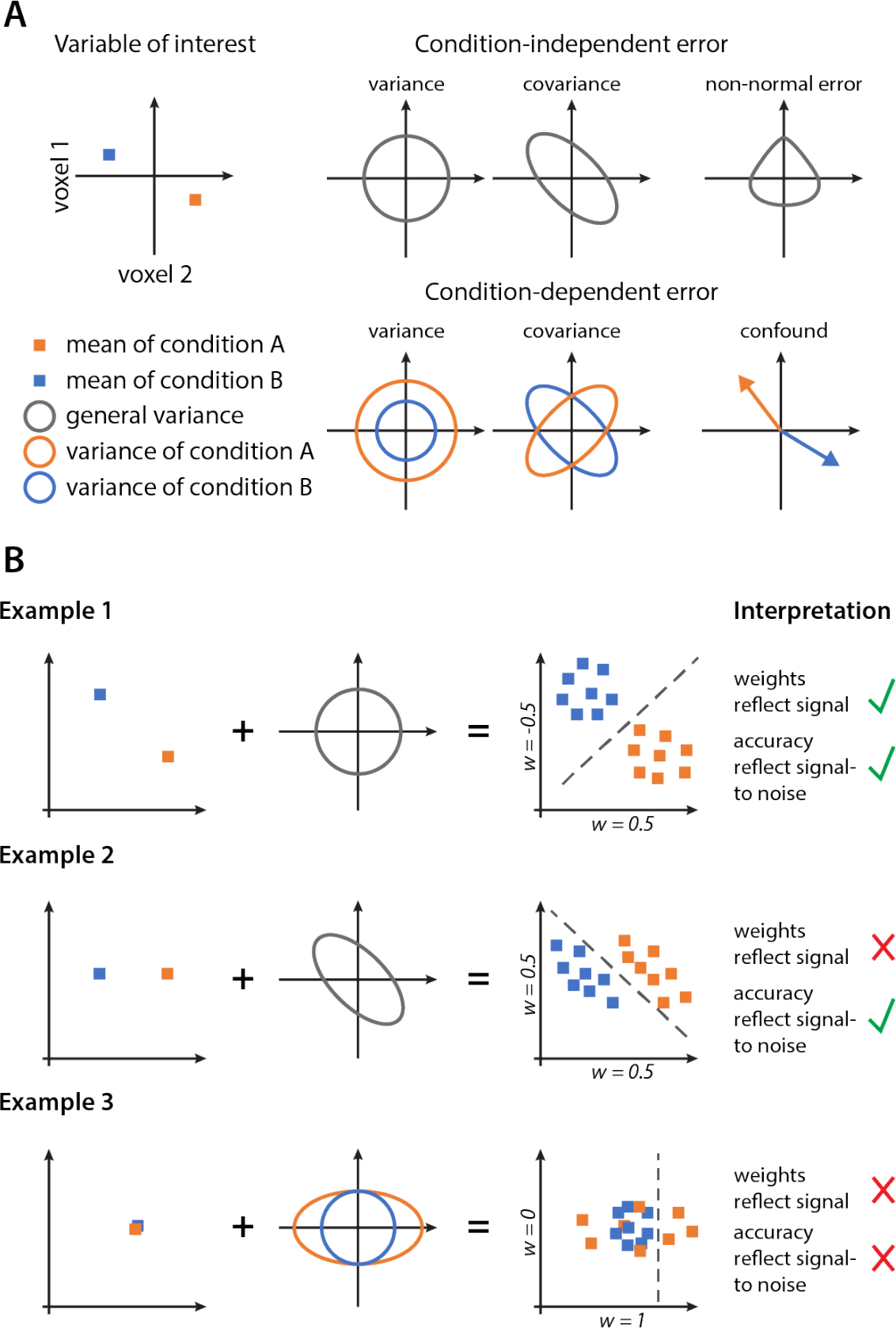
The prevailing view of signal and noise in neuroimaging, and its correspondence to information content in multivariate decoding. A. Motivated by the activation-based philosophy, signal reflects the multivariate means of the data, while noise can be either condition-independent error (variance, covariance, or non-normal error), or condition-dependent error (heteroskedastic variance or covariance, or confounds correlating with the conditions). B. Three examples comparing the correspondence of signal and signal-to noise with the weights and accuracy of a linear classifier. In Example 1, the classifier weights reflect the signal, and the accuracy mirrors the signal-to-noise ratio. In Example 2, noise covariance picked up by the classifier causes a departure from this correspondence. In Example 3, despite the absence of signal, differences in noise distribution allow above chance classification, leading to a non-correspondence of the signal to the classifier weights and accuracy.

Another important feature of this common activation-based view is that for two conditions, the *size of the difference* between the mean parameters reflects the *signal strength*, and the *ratio of this difference to the noise component* reflects the *signal-to-noise ratio*. In other words, one voxel is perceived as more activated when it has a larger parameter value than another voxel, and this difference in parameter values directly reflect the signal.

### The multivariate decoding view of signal and noise

In contrast to the activation-based view of multivariate patterns depicted above, in multivariate decoding the focus lies on what information about the experimental conditions can be extracted from the measured response. To avoid confusion about the terminology of signal and noise, here we use the term *information* to describe what is signal and noise in this methodological approach. For multivariate decoding studies that aim at the interpretation of activity patterns discussed above (*multivariate decoding for interpretation*), linear classifiers are the most common choice. They are commonly chosen, because they generally perform well (Cox and Savoy, 2003; Misaki et al., 2010), they don’t overfit as easily as nonlinear classifiers, their parameters are more easily interpretable, and they provide a plausible lower bound of the information that another brain region can potentially read out (Kriegeskorte and Bandettini, 2007; Naselaris et al., 2011). In linear classification, each voxel receives one weight parameter, and the product of the weight vector and the measured response pattern across voxels is used to assign class membership to that pattern. In that respect, a large absolute weight reflects a stronger contribution of that voxel to the final classification.

Since the goal of multivariate decoding is discrimination of the experimental conditions, any component of the measurement that contributes to their discrimination is information, while any component that does not affect or reduces discriminability is not. This definition has an important consequence: not only differences in the means, but also differences in the data distribution can be information for a classifier. Further, as has been pointed out recently (Haufe et al., 2014), even data covariance that alone does not allow discrimination between conditions can contribute to the classification by suppressing correlated noise in the response and improving classification. Even though this variability contributes to the discrimination, it is not a source of information because it alone does not allow discrimination. This will become clearer in the examples below.

In this information-based view, the signal-to-noise ratio translates to the predictive accuracy of a classifier. Importantly, a weight parameter does not reflect the discriminability of each voxel in isolation. Instead, the absolute value of a voxel’s weight parameter directly reflects the *usefulness of that voxel considered as the contribution to the discrimination process* in the context of the other voxels included in the classification analysis.

### The collision of signal, noise, and information

To illustrate how this view of signal and noise impacts our interpretation of data and results, we will consider three examples (Figure 3B). In these examples, the data generation process follows the standard statistical framework, described as a linear combination of signal and noise components. Once the data is generated from these components, a linear classifier is applied to classify this data: It assigns weights to each of the voxels and measures information content based on these data. In each example, we assess two properties: First, do the weights of the classifier also reflect signal strength? Second, does the classification accuracy also reflect the signal-to-noise ratio?

### Example 1: Signal plus zero covariance Gaussian noise

In this first example, the measurement is described as a combination of a signal component and Gaussian noise with no covariance. A classifier could now read out this information by appropriately combining the two sources of signal. Since there is no covariance and the errors are Gaussian, it has been shown that the best classifier in this context is a Gaussian Naïve Bayes classifier (Zhang, 2005). The classifier places weights based on how much signal there is in each voxel, i.e. the weights reflect the signal strength in each voxel. In this case, the classification accuracy will closely reflect the signal-to-noise ratio.

### Example 2: Signal plus Gaussian noise with covariance

In this second example, the measurement consists of a combination of a signal component, where only voxel 2 distinguishes the two classes, and Gaussian noise that exhibits negative covariance between voxels, i.e. when one voxel’s response increases, the other voxel’s response will decrease. In this case, the Bayes-optimal classifier is the Fisher linear discriminant (Bishop, 2006). Importantly, the weights still represent how useful each voxel is for the discrimination of the classes; however, the weights no longer reflect the signal strength but a combination of signal and noise. The classification accuracy on the other hand still reflects the signal-to-noise ratio of the multivariate data.

### Example 3: No signal plus heteroskedastic Gaussian noise

In this third example, the measurement exhibits an absence of any signal and consists only of noise. In other words, the expected value of both conditions is the same. The noise exhibits no covariance. While the noise in voxel 1 has the same variance in both conditions, in voxel 2 it varies more strongly for the orange condition than the blue condition. A simple classifier such as a linear support vector machine can now separate the data points in a way that leads to above-chance classification: one condition is always classified correctly, while the other is only sometimes misclassified. Thus, there is information present that allows the discrimination of the classes, despite the absence of what we normally describe as signal. This is a property that holds for any linear classifier, because as soon as there is variability in the estimation of the hyperplane and a deviation of this hyperplane from the center of the distributions, there will be above-chance classification^18^. This property is not specific to using accuracy as an information estimate, but also occurs for other popular information estimates such as d-prime or area under the curve. Further, an optimal nonlinear classifier could easily provide a much higher classification accuracy. In this example, the weights do not reflect the signal strength of each voxel, but reflect the variability of noise. In addition, the accuracy does not reflect the signal-to-noise ratio: The variability in the measurements, which is treated as noise in the standard statistical framework, translates to information in the information-based framework (Görgen et al., this issue).

These three examples reveal an important but often underappreciated fact: Multivariate decoding depends not only on what we commonly treat as signal – differences in the multivariate means - but also on what we treat as noise – the variability of the measurements. This has three consequences. First, the weights of a linear classifier cannot be interpreted to reflect the signal, but only to reflect the importance of each voxel for the classification process (Haufe et al., 2014). Second, the information content measured with a classifier (e.g. prediction accuracy) not only reflects differences in multivariate means, but can also purely reflect differences in variability (Davis et al., 2014; Görgen et al., this issue). Third, for a classifier to generalize to unseen data, it not only requires stability in the signal, but also stability in those components of noise that contribute to the classification.

One may wonder what factors affect the noise covariance of the data and under what circumstances there would be different noise covariance between conditions that could translate to above-chance classification accuracies in the absence of “signal” (see *Example* 3). After all, if these differences were indeed of neural origin and reflected the variable of interest, this information could reflect a processing strategy employed by the brain. Thus, such results would demonstrate that methods in the information-based framework such as multivariate decoding are sensitive to information that would be missed by methods in the activation-based framework. Indeed, the study of noise covariance is growing in popularity in animal electrophysiology (Averbeck et al., 2006; Churchland et al., 2010; Ponce-Alvarez et al., 2013) and neuroimaging (Garrett et al., 2011; Kohn et al., 2009).

Central to this discussion, however, is whether the differences in noise covariance can meaningfully be attributed to i) neural variability and ii) the variables of interest. In fMRI, non-neural factors commonly affect noise correlations between voxels. These include physiological noise such as head motion and noise fluctuations related to the cardiac / respiratory cycle, and separating those from neural sources of variability is difficult as demonstrated in the analysis of functional connectivity (Power et al., 2016). Even if differences could meaningfully be attributed to neural variability, it needs to be determined that this variability is related to the condition of interest and not other uncontrolled confounds. Thus, many differences in noise covariance may not be specific to the variables of interest, but could be caused by other factors. As we will point out below, even the experimental design in the absence of data can induce differences in the variability of conditions. Thus in a classical decoding setting, it may turn out to be difficult to disentangle neural variability of interest from other sources of variability.

## 5. Interpretation of multivariate decoding

So far, we have laid out the differences between multivariate decoding for prediction and multivariate decoding for interpretation, described the differences between classical univariate analysis and multivariate decoding, and illustrated in the different interpretation of signal and noise in a standard statistical framework and the information-based framework. Here, we use four illustrative examples to highlight how these differences in frameworks may translate into confusions related to the interpretation of results using multivariate decoding. In particular, we focus on examples that demonstrate how the theoretical considerations described above may impact the application and interpretation of multivariate decoding for the study of brain function. Crucially, these examples do not invalidate the methods used. Rather, they are meant to highlight potential confusion regarding the motivation of these approaches, their interpretation, and what may happen when their assumptions are violated.

### 1. Interpretation of low decoding accuracies

In multivariate decoding for prediction, the goal is to build a classifier that can be used in real-world applications. In this approach, decoding accuracies that are close to chance indicate that the classifier is far from this goal, which questions the usefulness of this approach in practical applications, either because of data limitations or because of the chosen classifier^19^. Even though in multivariate decoding for interpretation the focus is not on real-world applications, it is not uncommon for researchers (and reviewers) to question low decoding accuracies. This may arise because decoding accuracy is equated with effect size, and low decoding accuracies are treated as an indication of a small effect. Consequently, a small effect could be interpreted to indicate that a variable does not play much of a role in that brain region.

While it is true that for a given analysis classification accuracy reflects the size of an effect, accuracy does not reflect a standardized measure of effect size such as Cohen’s d. As illustrated in Figure 4, the accuracy depends heavily on averaging carried out prior to decoding (Allefeld and Haynes, 2014; Mumford et al., 2012) or the cross-validation scheme used, to name only a few. Consequently, a high accuracy can reflect a small effect (Combrisson and Jerbi, 2015), and differences in accuracy need not reflect differences in effect size or statistical power (Ku et al., 2008). Indeed, even accuracies close to chance can carry useful information if they generalize across the population (Christophel et al., 2015)^20^. Similarly, accuracies are bound at 100 %, adding to the difficulty of directly linking accuracy to effect size. Finally, even if decoding accuracy reflected effect size, it is difficult to interpret accuracy as the importance of that variable in a brain region, because response patterns may be less distributed in one region as compared to another, affecting the read-out without reflecting the importance of that region. Thus, if any, accuracy only reflects a relative measure of effect size, either within a given study across comparable conditions, or between studies when manipulating individual processing choices (but see Bhandari et al., 2017). Unfortunately, there are no straightforward ways to attain standardized effect size estimates for multivariate decoding. For example, classical standardized effect size measures such as Cohen’s *d* are invariant to averaging by taking into account the number of measurements and their dependence structure (e.g. temporal autocorrelation). An equivalent way of correcting for the number of measurements while accounting for correlated measurements is difficult if not impossible in multivariate decoding. For that reason, until such methods have been developed, it is probably advisable not to use information estimates derived from multivariate decoding as a measure of effect size for the comparison between studies, unless those studies use the same approach for generating results.

**Figure 4.**
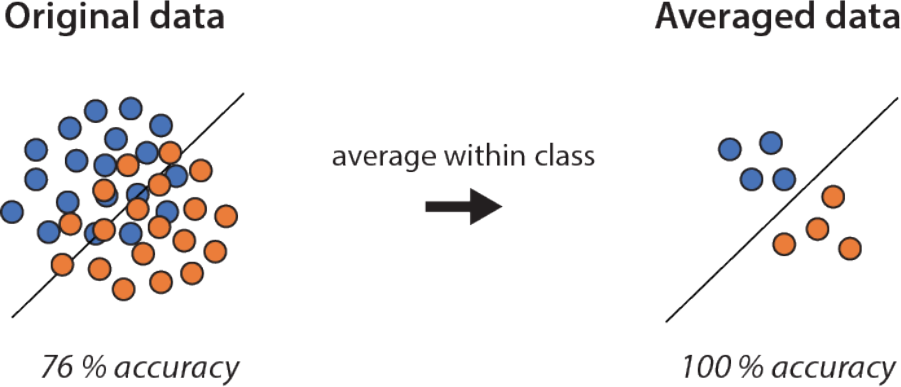
The accuracy of a classifier is not a standardized estimate of effect size, because it depends on choices such as averaging or the cross-validation scheme. For example, classification accuracies will be lower for single image decoding, but will increase when data within each class are averaged together. However, this need not translate to increased statistical power, because the accuracy estimate is based on fewer responses, increasing their variability. The confusion likely arises from the view that high decoding accuracies are necessary for a decoding model to be useful, which is often true in multivariate decoding for prediction but not multivariate decoding for interpretation.

### 2. Interpretation of univariate responses in multivariate decoding results

In many studies using multivariate decoding, researchers try to evaluate to what degree their results are reflecting univariate response differences between conditions. The motivation for interpreting univariate responses in the context of multivariate decoding varies. It might reflect the attempt to control for confounds that are assumed to lead only to univariate response differences (Coutanche, 2013), or to reveal multidimensional representations beyond “simple” one-dimensional activations (Davis et al., 2014). Alternatively, the motivation may reflect the idea that a “real” multivariate pattern is confined to subtle, fine-scale response differences and not mirrored in responses at a larger spatial scale accessible to classical univariate analysis (Freeman et al., 2011; Op de Beeck, 2010; Swisher et al., 2010). Finally, the motivation may simply be an effort to demonstrate the superiority of multivariate decoding. As we will see, and important to our discussion, the interpretation in fact does not reflect a comparison of univariate and multivariate responses, but what we described as uniform and non-uniform response differences.

One simple approach for getting at the difference in univariate and multivariate responses is comparing results of two analyses directly, for example by demonstrating a significant result with multivariate decoding but a null result with classical univariate analysis (for early studies, see e.g. Eger et al., 2008; Haynes et al., 2007; Kriegeskorte et al., 2007). A more common approach is to attempt removing univariate response differences between conditions from multivariate patterns (Jimura and Poldrack, 2012; LaRocque et al., 2013). However, it is unclear what is meant exactly be “removal of univariate response differences”, and what would constitute the “multivariate response” that remains after this removal.

In Figure 5, we depict three scenarios of what could be meant by removing a univariate response^21^. In the first scenario, the idea of removing univariate responses is interpreted as removing *any* univariate response differences between conditions from *every* voxel (Figure 5B). Since a multivariate response difference is based on univariate response differences, this removal would leave only noise variability as a basis for classification. Using a geometric interpretation with a space spanned by all voxels, this would correspond to the removal of the centroid of each condition in voxel space. While this is obviously not a realistic approach, it highlights the ambiguity of the term “univariate response” in the context of multivariate patterns.

**Figure 5.**
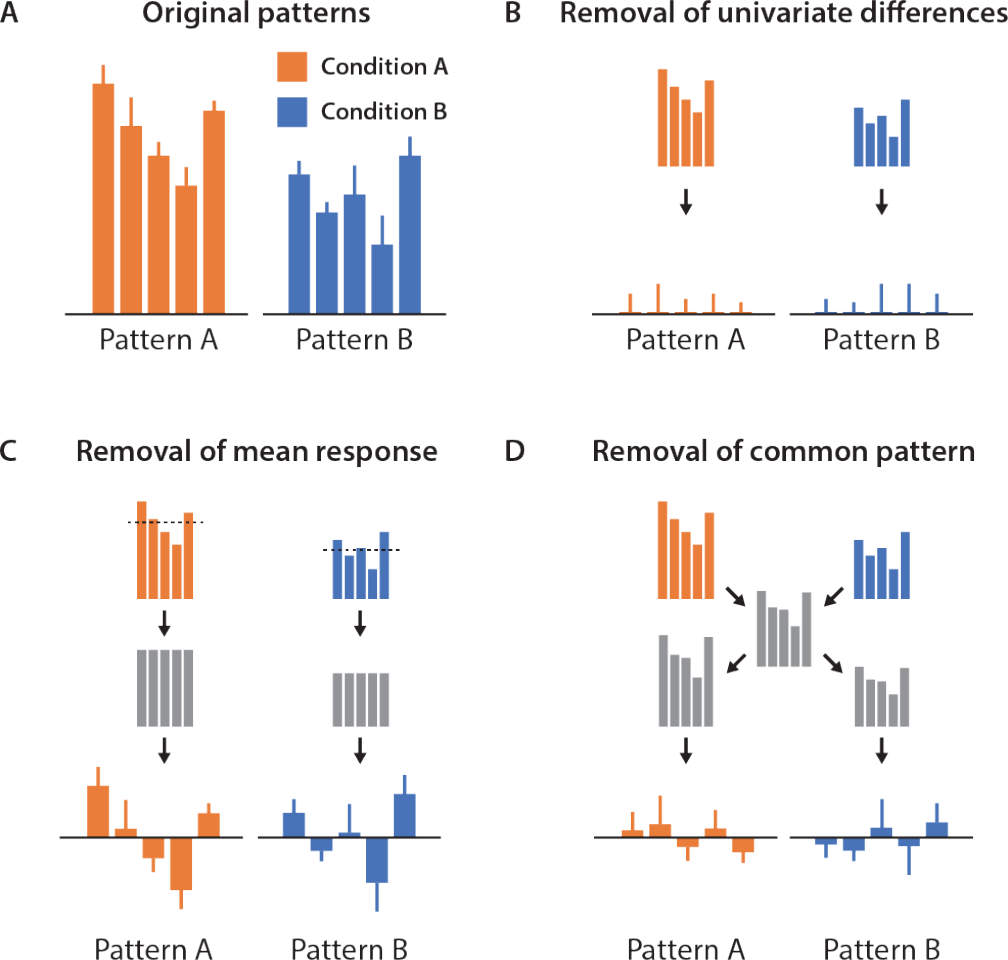
*Figure 5. Different interpretations of “removal of univariate response” from multivariate pattern. A. Original patterns. The response pattern is different across the two conditions. B. Removal of all univariate response differences. This approach removes any univariate differences between conditions from every voxel individually, leaving only the variability across trials. C. Removal of mean response. For each condition, the “overall activation difference” across voxels in a pattern is estimated and then removed from the response pattern. D. Removal of common pattern. The mean response pattern across both conditions is calculated and in another step scaled to optimally fit each individual response pattern. What remains as the corrected pattern is the (collinear) residuals of this fit.*

A second possibility is the removal of a uniform response across a pattern that is of the same sign and amplitude across all voxels, estimated as the mean response across voxels for each condition separately (Misaki et al., 2010, Figure 5C). This approach most closely matches the description of “overall activation differences” and is commonly employed in this context (Coutanche, 2013; Jimura and Poldrack, 2012). In the geometric interpretation, the univariate response corresponds to the projection of the data onto the (hyper)diagonal of voxel space, and the removal would shift the distribution of each condition along this diagonal towards 0. The approach assumes that the “univariate response” is identical in each voxel. However, this assumption is violated when there are differences in sensitivity between voxels (e.g. voxel 1 generally responds less than voxel 2), which, among others, may be caused by non-uniform distributions in neural selectivity, differences in neuronal density, differences in vasculature, or partial volume effects. When there are differences in sensitivity between voxels – which is almost always the case – this approach leads to incomplete removal of univariate response difference. In the geometric interpretation, the univariate response would no longer fall on the diagonal of voxel space, but for some voxels have a shallower angle when their sensitivity is lower than average, or a steeper angle when their sensitivity is higher than average.

Finally, the removal could refer to the subtraction of the common pattern shared between all conditions, which reflects a response that is of the same sign across voxels but allows for differences in sensitivity between voxels (Brouwer and Heeger, 2013, Figure 5D). This common pattern is estimated by first calculating the mean pattern across conditions and then fitting this pattern to each condition separately. In the geometric interpretation, this mean pattern would provide an estimate of the direction of the univariate response that no longer falls on the diagonal of voxel space, but is otherwise similar to the removal procedure described above. While this approach allows for a different amplitude in each voxel (Brouwer and Heeger, 2013), it assumes that the response pattern is only explained by this “univariate response”, an assumption that is violated as soon as there are additional responses that are not reflections of this univariate response. In the simplest case, this may be one or more voxels responding strongly irrespective of the condition. In the more complex case, this may be additional directions in the pattern that carry meaningful variance. Thus, this approach works only if the univariate response is sufficient to explain the measured response pattern.

Irrespective of the approach, the term “removal of a univariate response” falsely equates a multivariate response difference with a response difference that is of both positive and negative sign (a non-uniform response). However, as we have illustrated above, a multivariate response difference can have both uniform and non-uniform response components. This confusion likely arises because classical univariate analysis and multivariate decoding are contrasted directly, without distinguishing the multiple changes that occur when switching between the methods. While it is relatively simple to remove all univariate responses completely, the actual goal of removing the signed, uniform component of a response depends on assumptions. Thus, it is important i) to define what is meant by the removal of univariate responses, ii) to clarify the motivation for the removal and iii) to know the assumptions underlying this process. In many cases, signed response differences are a useful source of information to distinguish the categories of interest and can validly be included in the multivariate decoding analysis.

### 3. Interpretation of cross-classification accuracies

A popular approach in multivariate decoding is the use of cross-classification, which refers to the ability of a classifier to generalize between different contexts. As has been pointed out above, classification accuracy can be treated as a lower bound of the information content in a brain region. If a classifier trained on one context can generalize to data from another context, this demonstrates some degree of stability of the representation between both conditions and can be used to assess associations between cognitive processes (Kaplan et al., 2015). For example, a classifier trained on objects at one retinal position and tested at another can be used to test whether visual object representations are position-tolerant (Cichy et al., 2011; Kravitz et al., 2010). Likewise, a classifier trained on distinguishing items held in visual working memory can be used to test whether those items are represented similarly when they are the product of a mental rotation (Albers et al., 2013; Christophel et al., 2015). On neurophysiological data, it has become common to train a classifier at one point in time and test it at another to see whether it can generalize across time (King and Dehaene, 2014).

More recently, it has become common to interpret not only *whether* a classifier can generalize, but also *the degree to which* cross-classification is possible. For example, a representation may only be reported to be location-tolerant and not location-invariant, because the study demonstrated a decrease in cross-classification performance (Kravitz et al., 2010). Likewise, cross-classification in generalization across time is becoming more common to infer stable or dynamic representations (Stokes et al., 2013).

One assumption implicit to interpreting decreases in accuracies during cross-classification, however, is that a classifier is only sensitive to the signal and not to the noise in the data. However, as we have pointed out above, a classifier can utilize both signal and noise to carry out classification, and the classification accuracy depends on both. Consider the simple illustration in Figure 6A. Here the ability of a classifier to generalize depends on the noise level along the dimension relevant to the classifier. Consequently, the classification performance can be impaired when the classifier generalizes to a noisy dataset. To test whether cross-classification is affected by noise levels, it is possible to assess whether a classifier can extract information from the noisy dataset in the first place.

**Figure 6.**
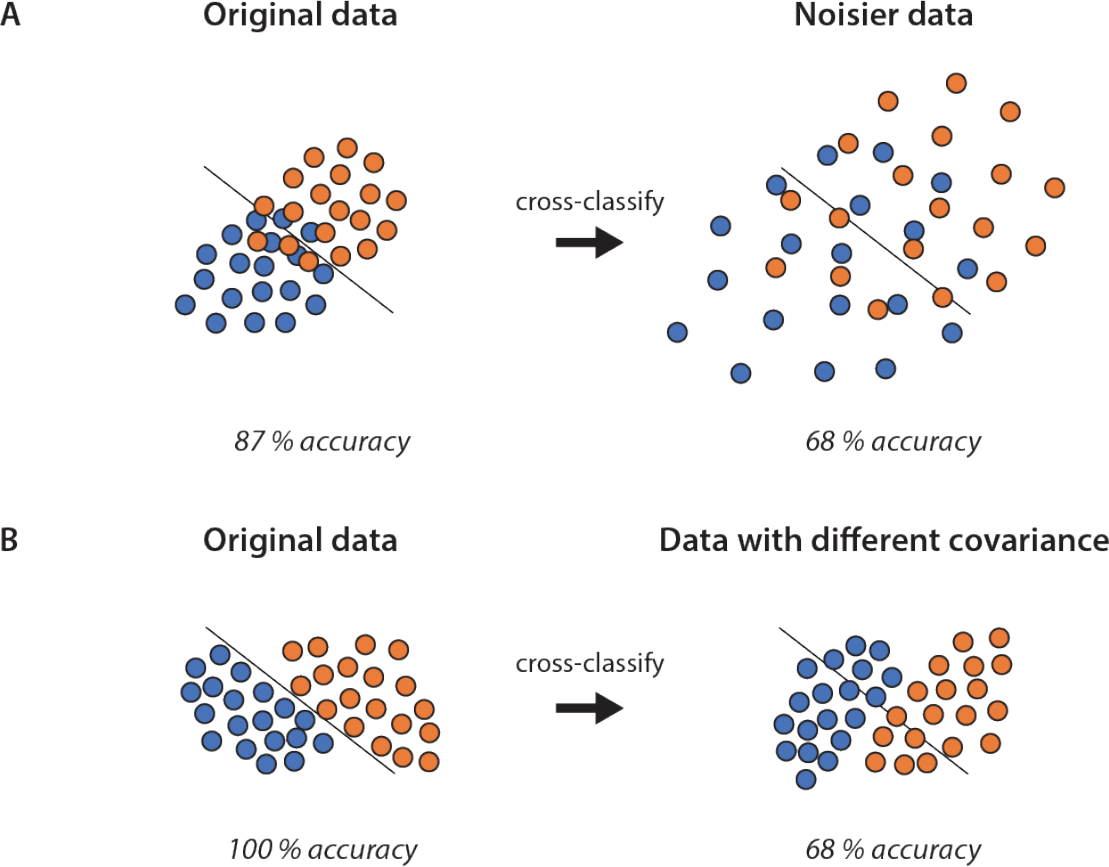
Effect of noise on cross-classification accuracies. A. Differences in the variability of data can affect cross-classification accuracies, despite there being the same effect in the difference of the multivariate means. However, a classifier trained on the noisy dataset would not perform well, either. B. Differences in the covariance of data can affect cross-classification accuracies, even when the general noise level does not vary. Here a classifier trained on the second dataset would perform equally, showing no asymmetries in classification or cross-classification.

A more complex example is shown in Figure 6B. Here, the classifier can distinguish both classes perfectly. However, cross-classification can be impaired even when the average response remains the same, but when the noise covariance is different between contexts. In a high-dimensional setting, this scenario depends on whether the direction of this covariance is relevant to the classifier, for example due to the presence of irrelevant brain responses that a classifier can filter out. Interestingly, in contrast to the previous example, here classification on the second dataset alone would reveal unimpaired decoding performance. The degree to which cross-classification is impacted by changes in the noise covariance depends on the intrinsic dimensionality of the data (Yourganov et al., 2011), which is typically much lower than the number of voxels. If the intrinsic dimensionality is high, it is unlikely for a classifier to utilize noise covariance and for changes in noise covariance to affect classification. This situation compares to the interpretation of weights described by Haufe and colleagues (2014), where noise covariance affects the weights of a classifier only if this covariance is used by the classifier to suppress noise. If the classifier is not affected by covariance in the data, the weights will more closely reflect the signal. Likewise for cross-classification, for data covariance not used by the classifier changes in the covariance will not affect the cross-classification performance.

Importantly, these examples do not invalidate the use of cross-classification. First, if cross-classification is possible, this demonstrates that signal and/or noise were sufficiently stable. Second, for cases where relative levels of cross-classification are interpreted, it is well possible that the assumption of stable noise is justified. Rather than discouraging the use of this method, our aim is to point out the assumptions underlying cross-classification, which may or may not matter in practice. Like the assumptions of a statistical test, it is useful to know how violations of a method’s assumptions can affect the interpretation of results.

### 4. Differential estimability of beta weights can lead to spurious decoding results

Multivariate decoding is commonly carried out on beta estimates from a GLM, which represent the conditions of interest. Beta estimates are often based on individual trials or the entire time-series, and different approaches have been suggested for their estimation in the context of multivariate decoding (Mumford et al., 2012). The estimability of a beta weight describes the expected variability of its estimation across many experiments. Among others, this estimability depends on the efficiency of the regressor, which can be calculated analytically (Dale, 1999). More variability in a regressor improves the estimability, and linear dependencies with other regressors reduce it. This has consequences for experimental designs in which the estimability is different between experimental conditions. For example, different number of trials entering each regressor can lead to differences in variability of the estimated beta weights, even in the absence of an effect (Görgen et al., this issue). Similarly, if the regressor of one condition exhibits a stronger linear dependence with the regressor of another condition, this affects the variability. In practice, this may happen for example when one condition is followed more often by a behavioral response than another, when one condition is more often preceded by a cue, or when stimulus jitter is not controlled appropriately. In Figure 3B, we described how a classifier can exploit differences in variability between conditions, despite the absence of differences in multivariate means. In the concrete example in Figure 7, this means that differences in estimability will lead to differences in classification, even when the data of both conditions come from the same distribution. That is, a classifier can perform above chance, because the estimability of the parameters in both conditions is different, not because there is a difference in the data. Importantly, this is an issue with the experimental design, not with the method used to attain pattern estimates^22^.

**Figure 7.**
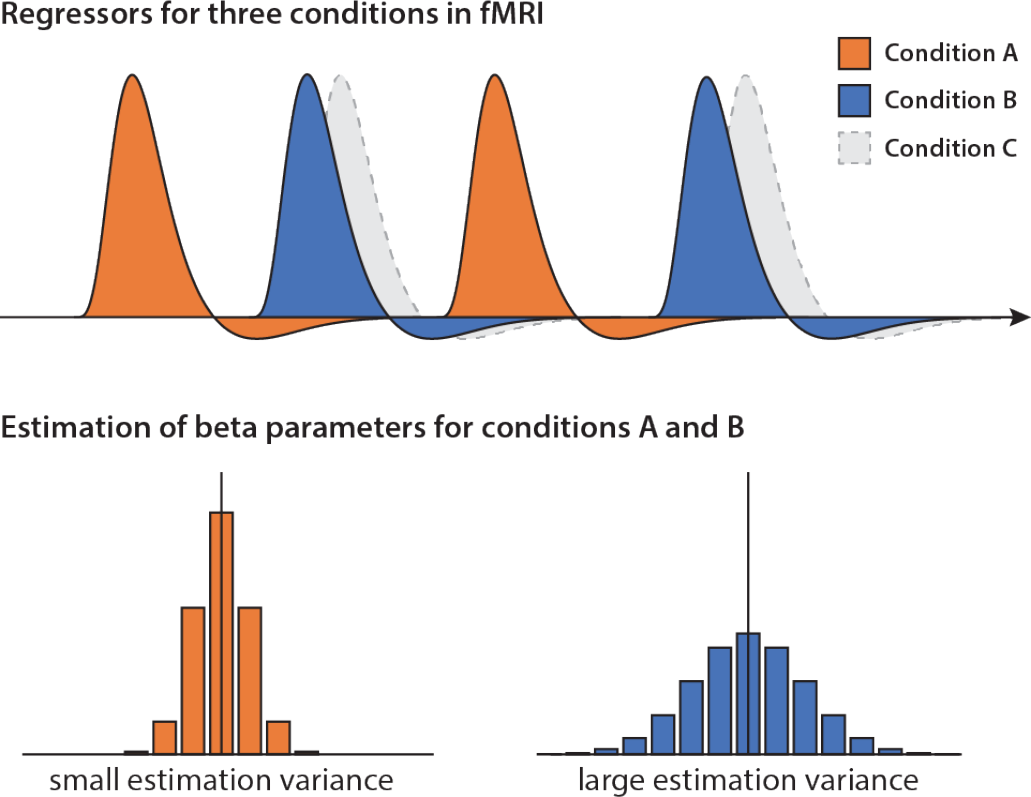
How differences in estimability between conditions can contribute to decodability despite an absence of differences in the data. The beta weights for Condition A can be estimated quite well, because this regressor is largely orthogonal to the other regressors, while the regressor for Condition B is non-orthogonal to the regressor of Condition C. As a consequence, on average, both beta estimates will be close to the true value. However, since the regressor for Condition B is non-orthogonal with Condition C, the estimation will be more variable. Classical methods would not reveal any differences between conditions. In contrast, as has been illustrated in Figure 3C, a multivariate classifier can pick up this difference in variability, which can lead to above-chance decoding accuracies even in the absence of any difference in the data. The reason for this discrepancy lies in the different meaning of signal and noise in the standard statistical framework and the information-based framework.

## 6. Strategies to resolve the confusions in multivariate decoding

In this article, we have described the current use of multivariate decoding for studying brain function and have highlighted confusions that arise from two issues. First, multivariate decoding was developed originally for making predictions and not for interpretations related to brain function. These different approaches, prediction and interpretation, have their own assumptions that may conflict with each other. Second, while multivariate decoding is embedded in an information-based philosophy, our thinking is still largely embedded in an activation-based philosophy, and we have demonstrated in this article that these philosophies are not always compatible. Further, the tools for statistical inference have been borrowed from the activation-based philosophy, adding to this confusion.

Moving forward, we suggest multiple strategies to resolve these confusions. Regarding the confusion of multivariate decoding for prediction vs. interpretation, we have two suggestions. First, we recommend researchers be more explicit about the goal of carrying out their multivariate decoding analysis. Is the goal building a predictive model that can serve as a biomarker for real-world applications, i.e. is the goal read-out of variables from the brain and maximal decodability? Or is the goal to learn more about the function of the brain? For a study of brain function, decodability in and of itself is not the goal; instead, the goal is what this decodability *implies.* Second, once this goal has been defined, we suggest researchers adapt their analysis specifically to this goal and not simply adopt existing dogmas in their analyses that may not apply to their goal. For example, as noted above, multivariate decoding for prediction necessitates high predictive value and out-of-sample prediction, but allows exploiting any consistent properties of the data. In contrast, multivariate decoding for interpretation does not require maximal prediction, but carries additional assumptions about what variables constitute signal and noise.

Regarding the confusion of multivariate decoding in the activation-based and information-based framework, we suggest two different strategies. First, when using multivariate decoding one approach is to carefully consider the assumptions that come with this approach and acknowledge the caveats this places on interpretation. As discussed above, these assumptions need not be limitations but can also expand our view of the representational architecture of the brain. Take the interpretation of the variability of measurements. On the one hand, successful decoding based on differences in variability may be perceived as an artifact, because information should only arise from signal, not from noise distributions. On the other hand, if this variability can be read out from a brain region, in principle it might also be used by another brain region as meaningful information. What matters in this context is whether differences in variability of measurements can be attributed meaningfully to neural variability, or whether they reflect other sources of noise that are unrelated to local changes in brain activity. In some cases, it may be difficult to know the assumptions and properties of a novel analysis strategy, despite us describing many properties of multivariate decoding in this article. In that case, we recommend the “Same Analysis Approach” that provides a principled approach to detect and avoid unanticipated properties of novel analysis methods (Görgen et al., this issue).

To limit the potential for confusion, a second strategy may be to employ alternative methods that increase sensitivity and specificity without requiring all the assumptions of an information-based philosophy, and that reduce the number of differences between classical univariate analysis and multivariate decoding. For example, cross-validated MANOVA (CV-MANOVA) is a powerful and versatile multivariate encoding method (Allefeld and Haynes, 2014) that provides cross-validated distance estimates that are estimates of the discriminability of variables of interest. CV-MANOVA is intimately related to the popular cross-validated Mahalanobis (crossnobis) distance estimate that is based on the linear discriminant (Walther et al., 2016). However, CV-MANOVA can directly be applied to time-series data, allows for estimating standardized effect sizes and provides all features of the linear model, including the use of multiple independent variables, the use of continuous variables, and the study of their interaction. Both CV-MANOVA and the crossnobis distance carry assumptions about signal and noise that are defined by the linear model, and using these methods the equivalent analysis for cross-classification does not suffer the interpretational difficulties discussed above. In the future, it may be possible to develop multivariate encoding approaches that allow researchers to choose between the study of uniform and non-uniform responses without cross-validation, which could prove fruitful when the focus lies on “overall response differences”. Researchers who are interested in the representational content of multidimensional representations or who want to test multiple competing representational models may use encoding models based on representational features derived from computational models, representational similarity analysis (Kriegeskorte et al., 2008), or pattern component modeling (Diedrichsen et al., 2011; Diedrichsen et al., 2017), the merits of which have been discussed in detail elsewhere (Diedrichsen and Kriegeskorte, 2017).

Having laid out the interpretational complexities of multivariate decoding, a critical reader may more generally question the usefulness of multivariate decoding for the study of brain function. Indeed, we believe alternative approaches for testing discriminability of brain measures, such as CV-MANOVA (Allefeld and Haynes, 2014) or the crossnobis distance estimate (Walther et al., 2016), may in many cases provide equal or higher sensitivity, while being more explicit about the assumptions, closer to our intuitions of signal and noise, and thus suffer from fewer interpretational difficulties. Both approaches are freely available in published software packages, (e.g. Allefeld and Haynes, 2014; Hebart et al., 2014a; Nili et al., 2014), making it easy to adopt them in research practice. Therefore, we think that in many cases researchers may want to consider departing from the use of multivariate decoding and use multivariate encoding methods instead. This switch would have the additional advantage of perhaps reducing the false sense of certainty that multivariate decoding offers direct measures of representational content, rather than being subject to similar interpretational ambiguities as standard statistical methods (Ritchie et al., 2017).

It is, however, worth noting that multivariate decoding for studying brain function has unique merits. It is sensitive to differences in the distributions of the data that multivariate encoding methods are not always sensitive to, unless modeled explicitly. In addition, some have suggested that, under certain circumstances and in conjunction with encoding methods, it is possible to use decoding to draw causal inferences about brain representations (Weichwald et al., 2015). Therefore, the choice of using multivariate decoding or switching to alternative methods should depend on the goal of the analysis (multivariate decoding for prediction vs. multivariate decoding for interpretation), on whether a researcher prefers a method with more explicit assumptions, and on the performance of the method in practice.

In summary, we believe that the use of multivariate decoding for interpretation can provide unique and valuable insights into brain function. We hope that our discussion of multivariate decoding helps clarify its role as an analysis method in the neurosciences, and that it aids recognition of the proper limitations and assumptions of this method in the study of brain function.

## Acknowledgements

The authors would like to thank Carsten Allefeld, Avniel Ghuman, Kai Görgen, Dave Jangraw, Daniel Janini, Niko Kriegeskorte, and Zvi Roth for helpful discussions and/or feedback on earlier versions of the manuscript. This work was supported by the Intramural Research Program of the National Institutes of Mental Health (ZIA-MH-002909) and a Feodor-Lynen fellowship of the Humboldt Foundation to M.N.H.

## Footnotes

1 For the reader unfamiliar with multivariate decoding, we provide a brief working definition. Multivariate decoding refers to techniques that jointly analyze multiple measurement channels (e.g. fMRI voxels) to make predictions about variables of interest. For categorical predicted variables, this approach reflects multivariate classification, while for continuous variables it reflects multivariate regression. Multivariate decoding is typically performed using machine learning algorithms, for example support vector machines. One instance of measurements across channels is described as a “pattern” (e.g. a multi-voxel pattern).

2 In the following, we use the terms “experimental condition”, “experimental variable” or “independent variable” not in the narrow sense as variables under the experimenter’s control (e.g. stimulus A vs. stimulus B), but in a broader sense including so called “quasi-experimental” settings, where the variable is under the environment’s control and selected post-hoc by the experimenter (e.g. participant’s choice A vs. choice B).

3 Competition website:http://www.lrdc.pitt.edu/ebc/2006/compoverview.htm, call for submissions:https://afni.nimh.nih.gov/afni/community/board/read.php?1,51415

4 The term *prediction* can have different meanings depending on the context. In inferential statistics, it refers to the existence of a model that can be used to tell how a variable will change in the future. For that reason, any model that describes a statistical dependence between two sets of variables can also be used as a predictive model. In the context of this article, prediction refers to models that are designed with a direct application in mind (such as stock market prediction), and where the reasons for this statistical dependence are only of secondary interest. While not irrelevant, space constraints preclude a discussion of the distinction between predictive models that allow predictions of dependent variables given the data without an explicit data generation model, and generative models that additionally allow making predictions about the data given the model (Bzdok, 2016; Naselaris et al., 2011).

5 Knowledge about the source of the information can help during the development of a new predictive model, when it is not yet clear if this source will help generalizing to all relevant cases. Using our example of the Pittsburgh brain interpretation competition, a non-neural source of information can and should be used for predictions if it is present in all relevant datasets.

6 Our use of the term *information* follows the common use in human neurosciences employing multivariate decoding, i.e. the presence of a statistical dependence in the data that can be read out with the help of machine learning methods and that is believed to be of neuronal origin. This use of the term does not imply that the brain region can communicate this information to another brain region or that it is used in behavior (Williams et al., 2007; De Wit, 2016).

7 One example of this is non-independence of training and test data, which would violate the assumptions of the prediction approach, but which may still allow meaningful inferences for interpretation when non-independence is modeled appropriately (Rosenblatt and Benjamini, 2014).

8 This interpretation is often causal, which in the absence of alternative explanations is a valid interpretation (Weichwald et al., 2015).

9 Others have discussed the parallel history of standard statistics and machine learning and how they differ (Bzdok, 2016). Here, the focus lies on the difference between activation-based and information-based philosophies and how they affect our interpretation of neuroimaging results. In our description, machine learning is just one methodological approach in the information-based philosophy.

10 It is important to mention that the two frameworks are not mutually exclusive, i.e. in principle they can measure the same statistical dependence and can both be restricted to the same types of relationships. For example, it is possible to convert some estimates from the standard statistical framework to an estimate of mutual information, and the Kullback-Leibler divergence that originated in information theory is common in frequentist and Bayesian statistics to estimate the difference between distributions. Despite this overlap, however, both frameworks nevertheless originate from different interpretational philosophies.

11 Here the terms “sensitivity” and “specificity” are not used in the classification sense of true positive and true negative response proportions, but to describe the discriminability and identifiability of variables, respectively.

12 Note that outside of neuroimaging, multivariate analysis is sometimes defined as the joint analysis of multiple *outcome* variables. However, in neuroimaging multivariate decoding typically has only one outcome variable, the experimental variable, and multivariate decoding refers to the prediction of that experimental variable by jointly analyzing multiple *measured* variables, typically measurement channels such as fMRI voxels.

13 Note that, while the difference in directional vs. non-directional analysis is closely related to uniform vs. nonuniform responses, both a uniform and non-uniform response can be analyzed in a directional and non-directional manner. For example, a directional analysis could reflect the pattern *difference,* while a non-directional analysis could reflect the absolute *distance* between patterns, a distinction that can be drawn for both uniform and nonuniform responses.

14 In the neuroimaging community, the term *encoding model* is often used in a narrower sense. In this narrower sense, first a computational model is used to mimic an alleged brain process. Then, it is tested whether the outputs of this model - typically representational features - are found to be encoded in brain activity. In this article, the term *encoding* is used in its more general sense, where any model is an encoding model that studies how a variable of interest is encoded in fMRI data.

15 It is not uncommon to interpret the parameters of a multivariate decoding model (e.g. the weight vector of a classifier) or to run statistical tests on them (e.g. Mourão-Miranda et al., 2005). However, these are neither activation estimates nor information estimates, as discussed below (see Haufe et al., 2014).

16 This only works if the multivariate approach does not always perfectly explain data (the upper limit is known as the capacity of an approach). For example, for linear classifiers in high-dimensional settings it is not unusual to reach perfect classification on the training data, which would likely not reveal any differences between iterations of a permutation test. Alternative unbounded measures of information content, such as the use of discriminative values or classical multivariate test statistics (Kriegeskorte et al., 2006), can circumvent this issue.

17 Our use of the terms “signal” and “noise” could alternatively be described as “components of the measurement that are of interest” and “components of the measurement that are not of interest”, respectively. While the terms are used inconsistently in neuroimaging (e.g. “brain signal”, “temporal signal-to-noise ratio”, etc.), we use these terms as a shortcut for describing relevant and irrelevant aspects of the measurements, which is close to their common use in cognitive neuroscience.

18 Note that, while this property holds for linear discriminant analysis (LDA) it does not apply to the closely related cross-validated Mahalanobis distance estimator, (Walther et al., 2016) which is an encoding method. While the accuracy of the LDA will increase with increasing differences in the variance, the cross-validated Mahalanobis distance estimator will on average remain the same but will become more variable.

19 There are exceptions, such as stock-market prediction where even a very low prediction accuracy can have enormous predictive value.

20 It may be argued that the actual reason for rejecting low accuracies is not effect size itself, but the idea that reported findings reflect a general positive classifier bias, either because of the classifier itself or because of the noise structure of the data. Indeed, this has led some researchers to the idea of estimating “empirical chance levels” using permutation approaches. Importantly, this caveat applies equally regardless of the accuracy level. When the analysis is free of nonindependence, then a bias reflects an uncontrolled confounding variable (Gorgen et al., this issue), and permutation approaches cannot easily deal with these cases.

21 Our discussion does not include the removal of the mean pattern, i.e. the mean response across conditions in each voxel. The consequences of this approach – also known as the cocktail blank – have been discussed elsewhere (Diedrichsen et al., 2011; Garrido et al., 2013; Walther et al., 2016). We did not include this approach, because the goal of this approach usually is not to remove condition-specific “univariate responses”, but to remove a pattern that is shared between all conditions. While this is similar to the approach described in Figure 5D (without additional scaling), it is not the motivation of this approach to completely remove univariate responses.

22 Note that this effect is different than a recently described bias in representational similarity analysis that occurs when using collinear regressors (Cai et al., 2016), because it more generally refers to the estimability of regressors, rather than only to their collinearity. While in principle it may be possible to at least correct for bias induced by collinear regressors by using the parameter estimate covariance matrix, this still needs to be demonstrated in practice and is expected to work less well under low signal-to-noise regimes. In contrast, the multivariate encoding methods described below do not lead to biased estimates.

